# Male-biased triploidy in *Populus tremuloides* and genomic structure of its southern relict populations

**DOI:** 10.64898/2026.01.12.695788

**Authors:** Roos Goessen, Chen Ding, Earl Raley, Lisa Tischenko, Jerritt Nunneley, Matthias Fladung, Christian Wehenkel, Nathalie Isabel, Ilga Porth

**Author notes:** retired (Matthias Fladung). <u>Authors for correspondence:</u> Prof. Ilga M. Porth, Dr. Roos Goessen.

## Abstract

**Premise:** Isolated populations at the southern range edge of aspen (*Populus tremuloides* Michx.) offer a unique opportunity to study ploidy, clonality and distribution of sex under arid, high-elevation conditions. We aimed to characterize such traits, including potential sex biases and associations with ploidy, investigating previously unexamined Texan populations in a range-wide genomic framework of genetic structure and demographic history of this keystone species.

**Methods:** We combined new genotypic data from western Texas (Davis Mountains, Big Bend, Guadalupe Mountains) with preexisting datasets, assessing range-wide genetic diversity, clonality, ploidy, and sex. We further conducted ADMIXTURE and phylogenetic analyses, reconstructed historical effective population sizes (*Ne*) with Stairway Plot 2.

**Results:** Texan stands revealed high clonality, with diploid and triploid genets. Most Davis and Guadalupe Mountain individuals clustered with a southwestern U.S. lineage of aspen, whereas Big Bend individuals grouped with a Mexican, demonstrating for the first time the continuous northward geographic distributions of allele frequencies in aspen coherent with phylogenetics. The significant male bias in sex ratios, particularly among triploids, suggests dimorphism in survival or reproduction. No effect of elevation on sex was identified. Demographic inference indicated an ancient bottleneck (∼1–2 Mya) common to all six lineages, but more recent historical *Ne* trajectories differ between the northern and southern regions.

**Conclusions:** These findings shed light on how clonality, ploidy, and sex have interacted in shaping dioecious plant species, highlighting the importance of incorporating such information into projections of how climate change and habitat loss will affect their distribution, abundance, and the extinction risk of their marginal populations.

## Introduction

Isolated mountainous stands of quaking aspen (*Populus tremuloides* Michx.) occur throughout the southern edge of the species’ range, including in Mexico (Goessen et al., 2022, 2025; Hernández-Velasco et al., 2025) and Texas (Nunneley et al., 2014). These relict stands provide a unique insight into a species’ genetic diversity and adaptive potential (Hampe and Petit, 2005). At the same time, such marginal populations may be particularly vulnerable to climate change, highlighting the need for understanding their current genetic composition and their capacity for persistence into the future. Stoddard et al. (2024) reported increasing aspen mortality among young, low-elevation stands throughout the mountainous study area, with a failure of recruitment into mature size-classes.

Aspen can reproduce both sexually, through seed, and asexually, via root suckers. In addition to diploids, triploid individuals are common, especially in the southern part of the species’ range, such as the western United States and Mexico (Mock et al., 2012; Goessen et al., 2022). In fact, many angiosperm species consist of multiple cytotypes with different physiological tolerances (Maherali et al., 2009). Triploidy in aspen has been linked to arid environments and may result from factors such as greater triploid viability or the production of unreduced gametes (Mock et al., 2012; Goessen et al., 2022; Blonder, 2024). The formation of such autopolyploids in nature goes hand in hand with their vegetative alterations related to differences in their cell sizes (larger), epigenomic changes, phytohormonal imbalances (particularly, auxins/cytokinins) and adventitious root sprouting (Mráz and Šingliarová, 2025). In fact, polyploidy bridges cells to entire ecosystems functioning (Fox et al., 2020). Because aspen is per definition an ecological keystone species (Rogers et al., 2020), its polyploid cytotypes can be at the center of rapid adaptation to a changing environment (Fox et al., 2020). Thus, an in-depth assessment of the geographical patterns in the frequencies of polyploid trees is warrantied (Ræbild et al., 2024; Schley et al., 2025). While no cytotypes have been assessed to date in Texan aspen stands, polyploidy occurrence can be expected given the arid local climate. Large clones have previously been linked to triploidy in aspen (Dewoody et al., 2008; Mock et al., 2008); with the Pando, a triploid male clonal forest covering over 43 ha in size, widely publicized.

*Populus* species are dioecious, with males as the heterogametic sex (XY system) in most of the genus (Geraldes et al., 2015; Müller et al., 2020), while some poplars have a female heterogametic ZW system including *P. alba* and *P. qiongdaoensis* (Müller et al., 2020; Li et al., 2023). However, little is known about how male and female individuals are distributed across landscapes. A previous study from Sichuan province, China, on *Populus purdomii* and *Salix magnifica* growing in sympatry on altitudes of 2,000 to 2,600 m found contrasting reproductive investments linked to altitude and sex in these related genera (Lei et al., 2017). A previous *P. tremuloides* study in the Colorado Rockies revealed sex ratio differences over elevation gradients. Female clones are more dominant at lower elevations and male clones are more present at high elevations, attributable to sex-related mortality (Grant and Mitton, 1979). In general, the relationship between sex and environmental gradients remains poorly understood. Interestingly, no sexual dimorphism (difference in non-reproductive functional traits) has so far been identified for the Pacific Northwestern species *P. trichocarpa* (McKown et al., 2017) or the European aspen sister species *P. tremula* (Robinson et al., 2014).

Recent studies (Goessen et al., 2022, 2025; Hernández-Velasco et al., 2025) based on Genotyping-by-Sequencing (GBS) derived genomic data demonstrated that most sampled aspen stands in Mexico are individual clones. Nunneley et al. (2014) assessed the clonal and genetic structure of individuals in isolated stands in Davis Mountains Preserve (West Texas) using microsatellites. Their study identified that three out of ten stands are part of the same multi-locus microsatellite genotype, while all other stands are individual clones.

In this study, we combined range-wide GBS data from previous studies (Goessen et al. 2022, 2025) with novel sampled individuals from several isolated stands in Texas, Davis Mountains Preserve, Big Bend National Park and Guadalupe Mountains National Park, which are part of the sky islands of the Trans Pecos region. We aimed to address the following questions: (1) What are the characteristics of sex, clonality, ploidy and genetic diversity of these Texan aspen stands? (2) How does the inclusion of this new data inform the range-wide population structure and phylogenetic relationships, as well as the historical changes in population sizes within aspen? (3) Is there any sex-bias present in aspen regardless of ploidy or in case of different ploidy levels? (4) Does elevation influence sex occurrence over aspen’s entire distribution range?

## Material and Methods

### Sampling and genomic data preparation of the *P. tremuloides* sample collection

This study included a total of 2,174 samples, encompassing 190 newly sequenced *P. tremuloides* samples, and a large previous *P. tremuloides* sample collection (n=1,903 individuals, 110 populations) and *P. grandidentata* (n=81, to check for potential hybrids) from our previous studies (Goessen et al. 2022, 2025). In the current study, the leaves of 122 *P. tremuloides* trees were sampled from three locations in Texas, *i.e.* Big Bend National Park (two stands), Guadalupe Mountains National Park (one stand) and Davis Mountains State Park (nine stands), in 2022 (**Fig. 1**). For further analysis, stands were grouped into four major populations based on geographical distance. DNA was extracted from dried leaf material as described in Goessen et al. (2022). For sequencing, in total, 140 Texas samples (including nine triplicates) and 50 duplicate samples from the previous collection were included totaling 190 samples (100 ng DNA per sample). A triple digest Genotyping-by-Sequencing approach was used for library preparation (3D-GBS, with restriction enzymes PstI/NsiI/MspI) and subsequently sequenced on Illumina Novaseq S4 (one lane). SNPs were called with the STACKS pipeline (Catchen et al. 2013) as described in Goessen et al. (2025). In total, 1,733,444 raw SNPs were obtained after SNP-calling. An overview of all samples is presented in **Dataset S1**.

**Figure 1.**
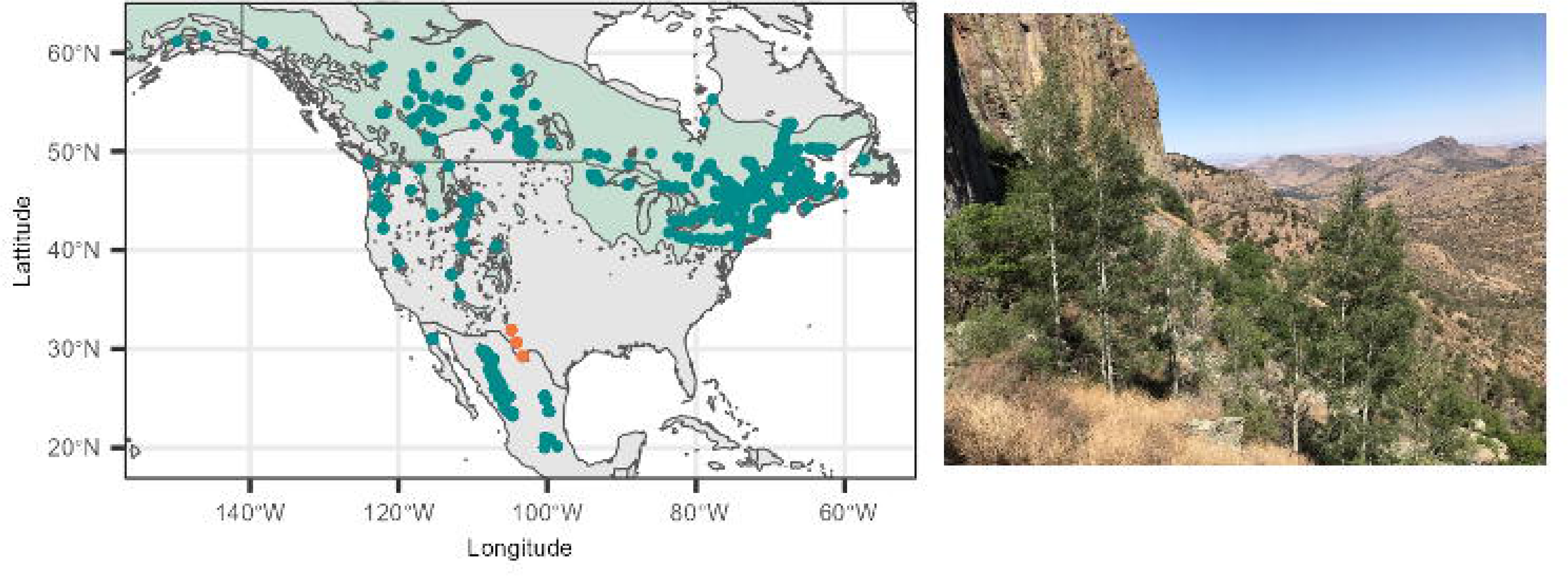
Individual aspen sampling distribution prior to filtering with photo of the aspen sampling effort at Laura’s Rock, Davis Mountains, Texas (photo: J. Nunneley, E. Raley). Dark turquoise dots are the included sampling locations from previous studies, and orange dots represent the locations of novel samples from Texas. Light green underlay represents the natural distribution of *P. tremuloides* (Little 1971). Respectively, one, nine and two natural aspen stands were sampled in Guadalupe Mountains National Park, Davis Mountains Preserve and Big Bend National Park.

### Ploidy and clonality assessment in Texan *P. tremuloides* individuals

We assessed clonality and ploidy among samples from Texas only, as the remaining sample collection has already been investigated (Goessen et al., 2022, 2025). To obtain many SNPs for ploidy assessment, we first filtered the raw VCF with loose settings using vcftools (––maf 0.05 ––max-missing 0.5 ––min-meanDP 12 –max-meanDP 100) (Danecek et al. 2011), which resulted in 77,836 filtered SNPs. We used FastPloidy (Goessen et al. 2022) to distinguish between diploid and triploid genotypes. We hereafter detected clonemate pairs using the shared heterozygosity index (SH), and samples with an SH above 0.9 were considered clonemates (Yu et al., 2023). We also filtered the dataset more strictly for clone assessment using the script available at https://github.com/enormandeau/stacks_workflow with parameters 8 90 0 68, without the relatedness step (see below for further details). We ran the script on both, the loosely filtered SNPs as well as the strictly filtered SNPs with both cytotypes included, since several samples were filtered out in the strictly filtered dataset due to missingness; in particular, the two stands DMP-06 and DMP-07 from Davis Mountains showed high missingness values.

### SNP filtering for further analyses of the *P. tremuloides* genomic dataset

For subsequent analyses, we filtered the entire dataset more elaborately; we removed *P. grandidentata* samples (see Goessen et al. 2022), removed duplicates, and made separate datasets with only diploid individuals and mixed ploidy, respectively, and then filtered the raw SNPs following the script in Goessen et al. (2022), which is based on the original script developed by Eric Normandeau (https://github.com/enormandeau/stacks_workflow) with parameters 8 90 0 59 and 8 90 0 68 (respectively, for diploids and mixed ploidy), and includes amongst others a relatedness filtering step. These four preceding numbers correspond, respectively, to the minimum allele coverage to keep a genotype, the minimum percent of genotype data per population, the maximum number of populations that can fail the maximum missingness, and the minimum number of samples with rare allele (mas). The diploid dataset encompassed 891 individuals and 13,033 SNPs, while the mixed ploidy dataset included 938 individuals and 12,519 SNPs.

For clonality assessment over the entire sample size (see below, without Texas samples) we used the same filtering settings as described above for diploids (8 90 0 59) however without performing a relatedness filter, resulting in 1,549 individuals and 15,259 SNPs.

For demographic history analysis with Stairway Plot 2 (see below) we used only diploid samples and filtered for a mas of 3 over the entire range (thus using settings 8 90 0 3), as higher mas values can influence the demographic inference. This dataset resulted in 52,327 SNPs, and we kept only individuals that were also assessed for population structure (i.e. 891).

### Population structure, genetic diversity and phylogeny assessments on the entire *P. tremuloides* genomic dataset

We ran ADMIXTURE (Alexander et al., 2009) with 50-fold cross-validation for K=2 until K=10, to assess population structure and perform appropriate K-number selection. We used RAxML to assess species-wide maximum likelihood phylogeny (Stamatakis, 2014), using the GTRGAMMA model and 1,000 bootstrap replicates.

For the diploid dataset, diversity statistics (observed heterozygosity (*H*_OBS_), within-population gene diversities (*H*_S_) and inbreeding coefficient (*F*_IS_)) were calculated per lineage using the function *basic.stats()* from the R package *Hierfstat* (Goudet, 2005). We used the function pairwise. *WCfst()* from *Hierfstat* to calculate the pairwise Weir and Cockerham *F*_ST_ value between lineages (Weir and Cockerham, 1984).

### Demographic history analysis per *P. tremuloides* lineage

We ran the demography analysis on the genetic clusters identified by ADMIXTURE; samples were separated based on population means for the appropriate K values for diploids. We used EasySFS.py (https://github.com/isaacovercast/easySFS, version 26-04-2022) to convert our VCF file to a folded site frequency spectrum (SFS). We identified the following projection values for downsampling the dataset (based on highest number of segregating sites) for each identified genetic lineage: 370 (NW), 78 (SW), 90 (PW), 266 (WSM), 46 (ESM), 636 (NE). For explanation of the lineage abbreviations used, see Results section.

We employed Stairway Plot 2 to assess the historical demography of the species (Liu and Fu 2020) using the folded SFS. To assess the parameter “L” (number of sequences), we calculated the SNP density per 125 bp (the read length used by STACKS) using VCFtools, resulting in SNPs being distributed over 11068 intervals of 125 bp, and we thus used an L of 1383500 (11068 * 125). We ran the program with the singleton SNP masking option included, as Stairway Plot 2 is sensitive to singletons (Liu and Fu, 2020).

### Sex determination across the *P. tremuloides* sampling

Sex assessment was based on the presence/absence of the *TOZ-19* gene in aspen. This gene is male-specific and can be verified by PCR and agarose gel electrophoresis following an established approach (Pakull et al. 2015). In addition, a total of four panels of 38 biallelic SNP markers were developed by the Canadian Forest Service (Natural Resources Canada) using the Sequenom® iPLEX® Gold Genotyping MassARRAY Technology (Agena Bioscience, San Diego, California, USA). And while these panels encompass a total of 152 intraspecific variable SNP markers previously selected for detecting gene flow, they also include a SNP close to the diagnostic *TOZ-19* gene that was demonstrated to correctly infer sex.

The following methods were employed in assessing aspen samples for sex using the *TOZ-19* marker system: 21 Texas samples (representing all sampled populations there) at the IBIS institute (Quebec, Canada) using a newly developed multiplex PCR assay, 877 samples from Mexico at the Göttingen institute (Germany) and 530 samples from the rest of the aspen distribution range at the IBIS institute using the Pakull protocol. In addition, 388 samples were determined by Sequenom MassARRAY. Among all these samples, 31 were analyzed by both, via Sequenom and the Göttingen institute, and four were tested at the IBIS and Göttingen institutes for validations. Out of all validated samples, we report only one inconsistent sex assignment in aspen. We further removed 52 samples due to sampling bias in sex, where several female trees from Quebec and Utah were specifically intended for a seed germination experiment (Goessen et al. 2022).

To compare the sex, ploidy and clonality results of Texas samples to the entire aspen distribution range, we included the ploidy results from our previous study (Goessen et al 2025). Moreover, we performed clonality assessment using the SH-index (Yu et al 2023) with a cut-off of 0.9 over the entire sample size (calculated separately for the Texas samples) and compared this to results of Goessen et al. 2022 (who used a subset of samples present in this study), resulting in a highly similar clone assessment (see Dataset S1 in Goessen et al., 2022 vs **Dataset S1** for the current study).

We then combined all our sex, ploidy and clonality results and corrected the data to have only 1 individual per clone, except in cases were both males and females were present within a single detected clone. In total, 13 clones were found to have both males and females, of which 3 were triploids and 10 were diploids; these individuals were mostly detected in Mexico, except for two cases found in Canada. We removed individuals that had missing data for either ploidy, sex or clonality. Finally, we used a chi-square test to assess whether a sex-bias is observed over the entire sample size and between cytotypes. We used a chi-square goodness-of-fit test to test against a null hypothesis of equal sex distribution, as well as to assess if sex distribution differs within diploids and triploids. Because our sampling strategy was not consistent over the entire range due to lack of *a priori* information, we could not infer whether there are effects of sex or ploidy on clonality.

We aimed to further assess sex distribution within the geographic area of each genetic cluster. We therefore first had to assign individuals that lacked cluster information to the previously identified genetic clusters based on latitude and longitude locations For this, we used the get.knnx function from the r package FNN (Beygelzimer et al., 2024) that uses a k-nearest neighbour searching algorithm.

Elevation was deduced using the *get_elev_point()* function from the *Elevatr* R package (Hollister et al., 2023) using our longitude and latitude data. We assessed whether there is a relationship between sex (dependent) and elevation (independent) using a generalized linear model (GLM) with logit link function as sex is a binary variable, over the entire range as well as within each genetic lineage.

## Results

### Sex, clonality and ploidy of *P. tremuloides* individuals from Texas

Among all Texas samples from Big Bend National Park (BIBE), Guadalupe Mountains National Park (GUM) and Davis Mountains State Park (DMP), respectively, a total of 30 triploids, 91 diploids and one ambiguous sample were identified (**Fig. S1, Table 1**). All triploids were found at Fort Davis and included all individuals in stands DMP05 (all males), DMP06 (all males) and DMP08 (all females). Besides site DMP08, individuals from stands DMP01, DMP02, DMP03, DMP04, and DMP07 were also all females, while individuals from stands DMP09, BIBE01, BIBE02 and GUMO01 were all males, besides the ones reported at sites DMP05 and DMP06 (**Table 1**). Using the SH-index under loose SNP filtering settings, we found that samples at DMP05 and DMP06 had SH values ranging between 0.66 (DMP05-9 vs. DMP06-9) and 0.95 (DMP05-10 vs DMP06-1) (**Fig. 2A**, **Table 1, Dataset S2**). Using stricter SNP filtering (**Fig. 2B, Dataset S2**), only one sample namely DMP06-11 remained (other samples at DMP06 were filtered out due to higher missingness); however, this sample had an SH-index above 0.9 with all samples of DMP05. Individuals in stands DMP05 and DMP06 are thus likely highly related or even clonemates (ramets), also considering that they were all assigned the same sex (all males). The closest individuals between those stands are 75 meters apart physically, while the furthest are 185 meters apart. Using the stricter filter, samples from population DMP02, DMP03 and DMP04 showed an SH-index above 0.9 and are likely also clonemates (all females). All other stands with an SH-index above 0.9 among all present individuals within a stand are thus clonemates (**Fig. 2B)**. Within DMP01, one individual DMP01-4 showed an SH-index of 0.86 with other individuals in the same stand. While slightly below the 0.9 threshold, we still considered it a clonemate within the DMP01 stand (**Fig. 2B).** Ploidy, sex, and clonal groups among Texan genotypes are visualized in **Fig. 3**.

**Figure 2.**
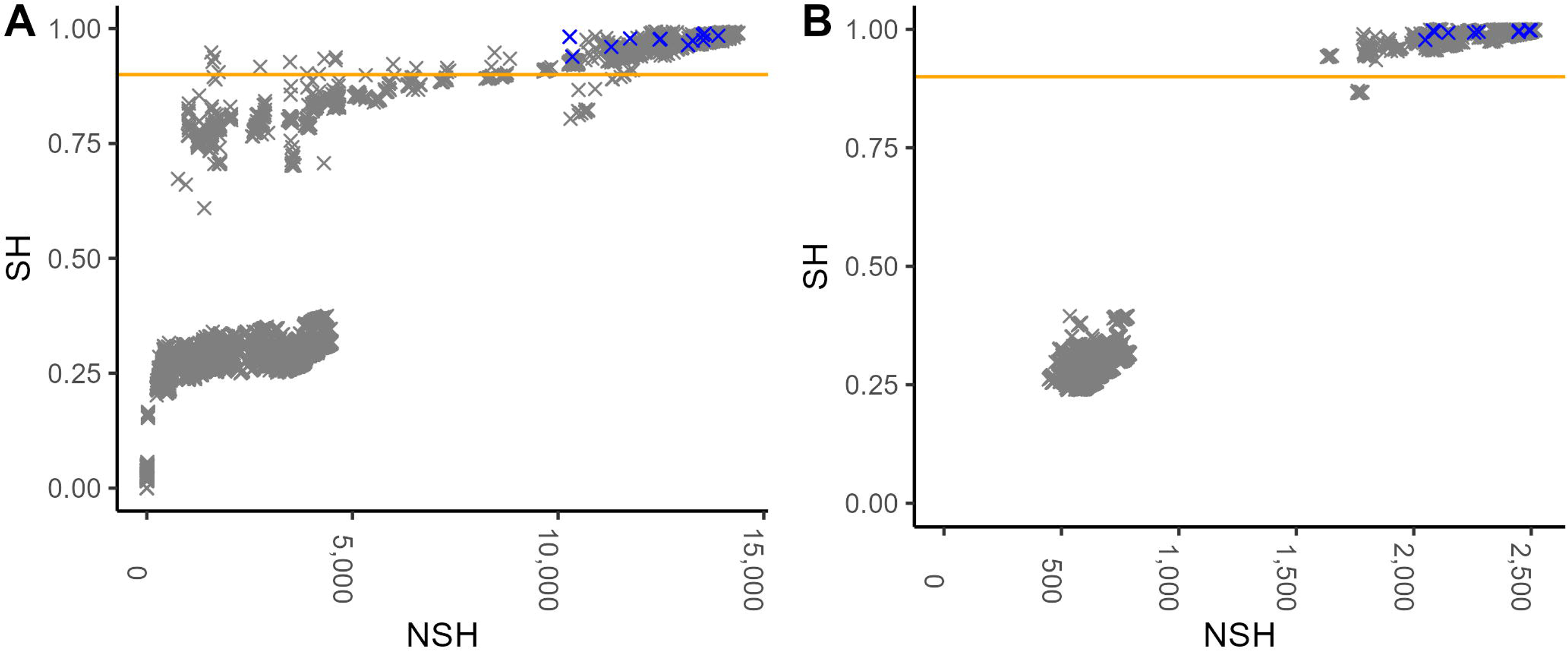
Shared heterozygosity (SH) index for all pairwise genotype combinations to assess clonality in Texan aspen. A. For the loosely filtered dataset. Samples #1-11 from DMP06, and samples from DMP05 have SH-index values ranging between 0.66 (DMP05-9 vs. DMP06-9) and 0.95 (DMP05-10 vs DMP06-1). **B. For the strictly filtered dataset.** Samples from populations DMP02, DMP03 and DMP04 have an SH-index > 0.9. Sample DMP06-11 has an SH-index > 0.9 with the present samples of DMP05 (#1-7). Here, all other samples of DMP06 and DMP05 were filtered out due to missingness. NSH is the number of shared heterozygous markers where two samples are identical. SH index represents NSH divided by the total number of heterozygous markers for the first sample in the pairwise comparison. Technical replicates are indicated in blue (n=11). The orange horizontal line indicates an SH of 0.9. See **Dataset S2** for detailed SH-index results.

**Figure 3.**
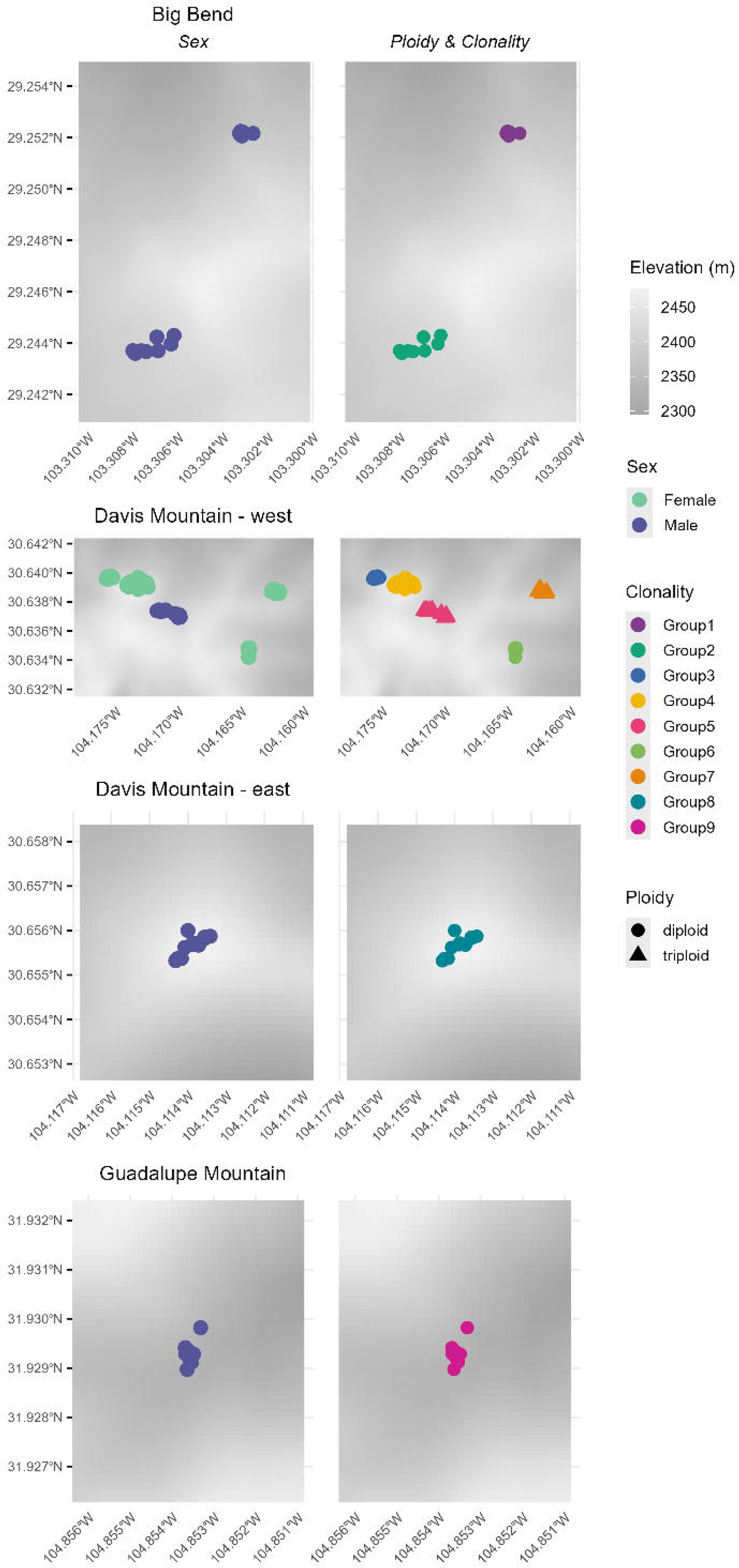
Clonality, sex and ploidy among Texan *Populus tremuloides* natural stands mapped onto their stand elevation profiles.

**Table 1.** Sex, clone number, and number of clonemates identified using the strict SNP filtering (SH-index above 0.85) or loose SNP filtering (SH-index above 0.6) per sampled Texan aspen stand. Indicated numbers do not include the technical replicates. Sex was determined on a subset of samples within each stand, with F meaning female and M male.

| Location | Stand | Sex | Size stand | Clone number | Clonemates_ strict (SH>0.85) | Clonemates_ loose filter (SH>0.6) |
| --- | --- | --- | --- | --- | --- | --- |
| Big Bend | BIBE01 | M | 10 | 1 | 10 | 10 |
| Big Bend | BIBE02 | M | 9 | 2 | 9 | 9 |
| Davis M. | DMP01 | F | 10 | 3 | 8 | 10 |
| Davis M. | DMP02 | F | 10 | 4 | 7 | 10 |
| Davis M. | DMP03 | F | 10 | 4 | 9 | 10 |
| Davis M. | DMP04 | F | 10 | 4 | 10 | 10 |
| Davis M. | DMP05 | M | 10 | 5 | 7 | 10 |
| Davis M. | DMP06 | M | 11 | 5 | 1 | 11 |
| Davis M. | DMP07 | F | 11 | 6 | 2 | 11 |
| Davis M. | DMP08 | F | 10 | 7 | 10 | 10 |
| Davis M. | DMP09 | M | 11 | 8 | 11 | 11 |
| Guadalupe M. | GUMO01 | M | 10 | 9 | 10 | 10 |

### Genetic diversity, structure and phylogeny in *P. tremuloides*

The lowest cross-validation value for the diploid dataset was K=6, while for the mixed ploidy dataset it was K=5 (**Fig. S2, Fig. 4**). In both datasets, two large northern clusters were identified, northeast north America (NE) and northwest north America (NW). In both the diploid and mixed ploidy data sets we further identified a Pacific western US (PW) lineage, and an inland southwestern US lineage (SW); the latter also included the populations from Baja California (Mexico) and the two most northern Texan populations (Guadalupe Mountains and Davis Mountains). For the diploid dataset (with K=6), within Mexico we observed the division between populations on the western Sierra Madre Occidental (WSM) and those on the eastern Sierra Madre Oriental (ESM). The mixed ploidy dataset (with K=5) found only a single genetic cluster within Mexico. Individuals from the Big Bend National Park clustered with the WSM genetic lineage. The phylogenetic tree revealed that individuals within genetic lineages are, as expected, phylogenetically close (**Fig. 5**).

**Figure 4.**
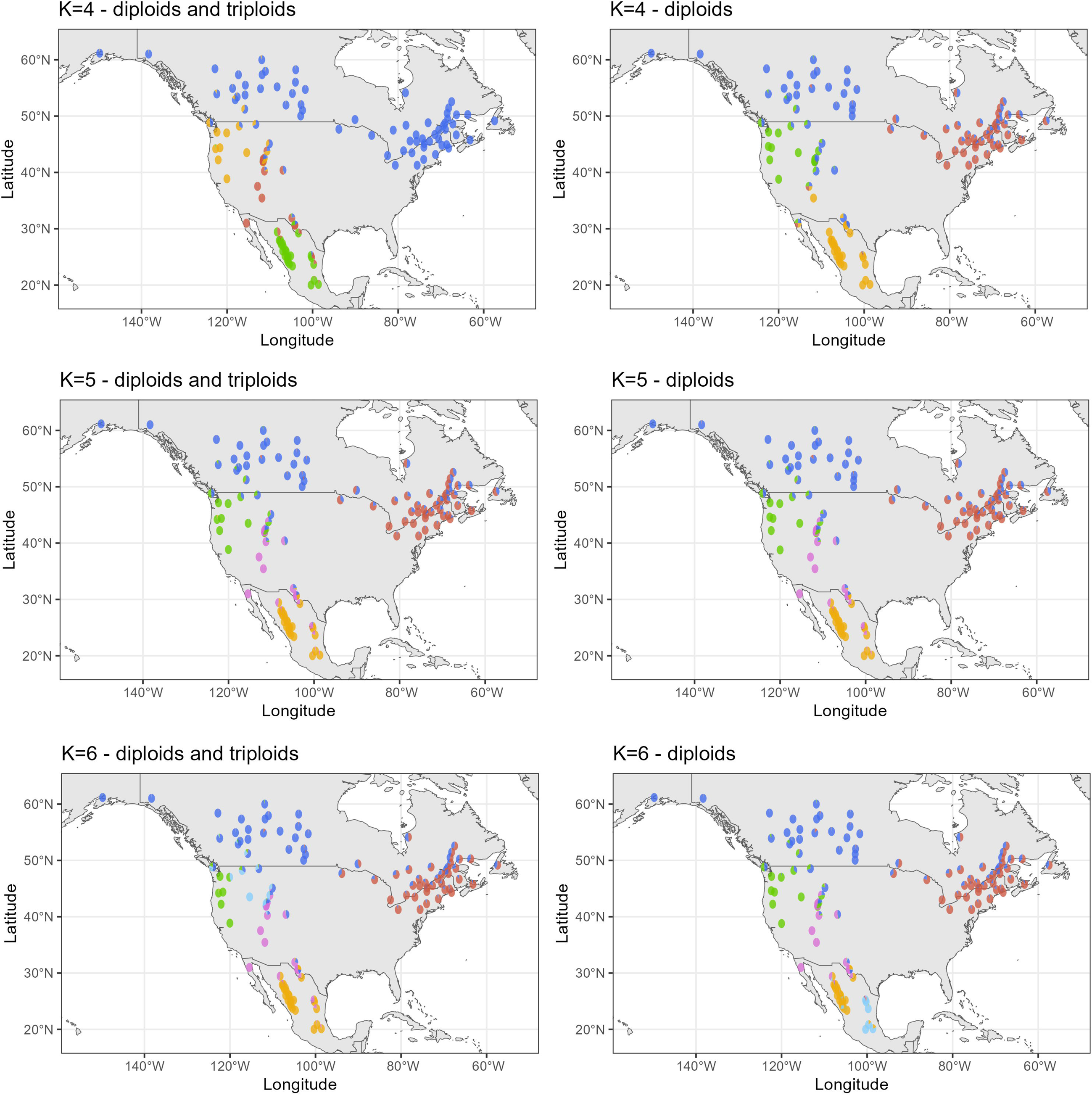
Admixture results for K=4, K=5 and K=6 for, respectively, diploids only and mixed ploidy, across populations of the sampled aspen distribution range. Using the lowest cross-validation value, the best K is K=5 for the mixed ploidy and K=6 for the diploid dataset (see **Fig. S2**).

**Figure 5.**
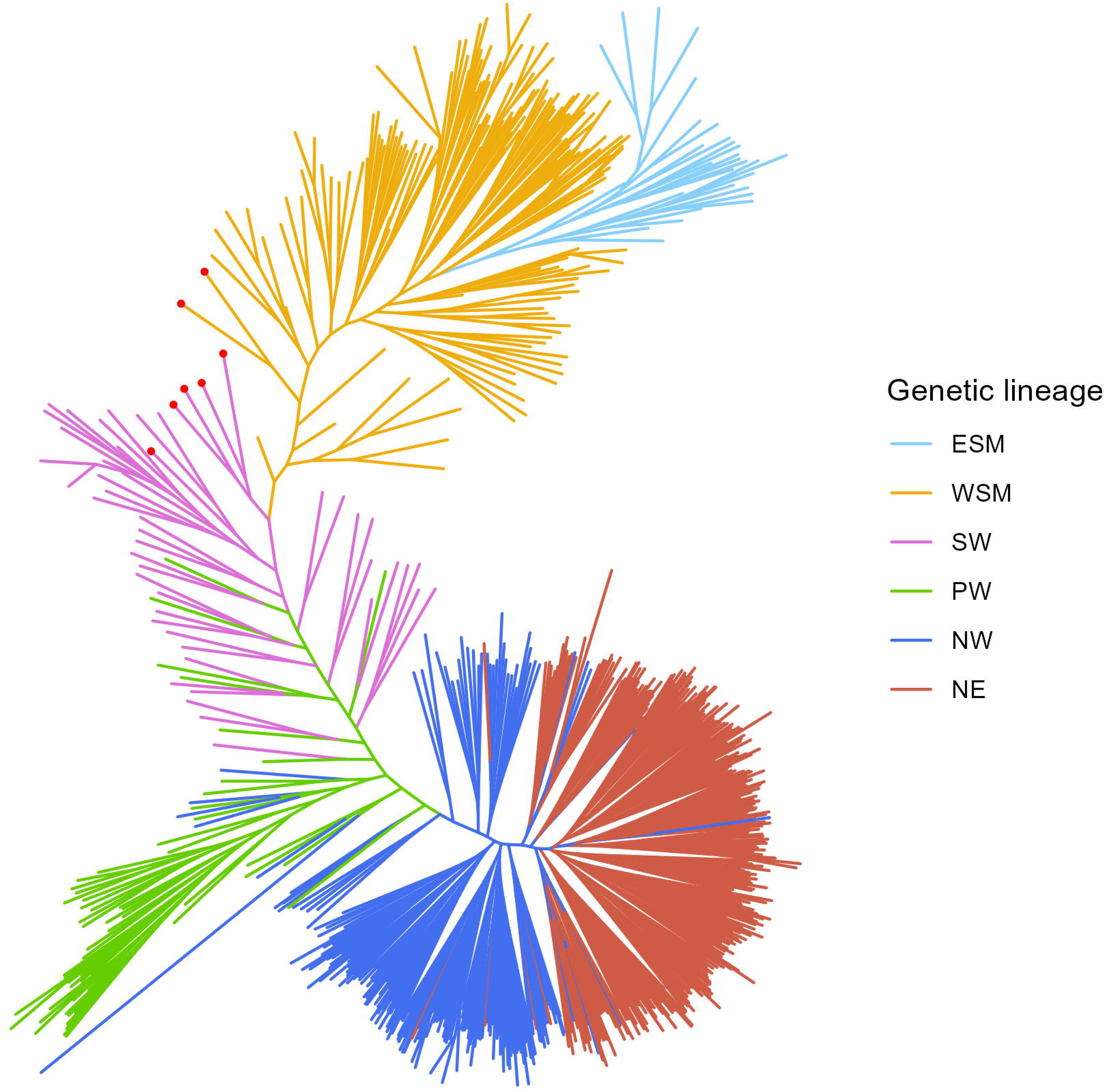
RAxML phylogeny for all individuals of the diploid dataset (K=6) across the sampled aspen distribution range. Genetic lineages are colored based on population means for admixture Q-values (See Fig. 4). ESM: Sierra Madre Oriental, WSM: Sierra Madre Occidental, SW: southwestern US, PW: Pacific West, NW: northwest North America and NE: northeast North America. Red dots indicate genets from Texas (7 in total).

The NE and NW genetic lineages are genetically least diverged (*F*_ST_ = 0.009), while the WSM and PW lineages display the highest genetic divergence with a pairwise *F*_ST_ of 0.207 (**Fig. S3**), which is surprising considering that WSM and NE genetic lineages are phylogenetically and geographically further apart (**Figs. 4, 5**). Still, overall high genetic divergence was observed between the Mexican lineages WSM and ESM vs. the PW, NE and NW lineages (pairwise *F*_ST_ values between 0.175 and 0.207, **Fig. S3**).

The NE and NW clusters displayed the highest *H*_OBS_ values of 0.204 and 0.206 respectively, followed by 0.198, 0.179, 0.176 and 0.174 respectively for clusters SW, WSM, PW and ESM (**Table 2**). The inbreeding coefficient was low overall, with the lowest *F*_IS_ for the WSM, NE and NW lineages (ranging between 0.05 and 0.06) and the highest *F*_IS_ within the PW cluster (0.11).

**Table 2.** Diversity statistics summarized per genetic lineage for the diploid dataset across the sampled aspen distribution range. With observed heterozygosity (*H*_OBS_), within-population gene diversities (*H*_S_) and inbreeding coefficients (*F*_IS_). ESM: Sierra Madre Oriental, WSM: Sierra Madre Occidental, PW: Pacific West, SW: southwestern US, NW: northwest North America and NE: northeast North America.

| Lineage | $H_{OBS}$ | $H_S$ | $F_{IS}$ |
| --- | --- | --- | --- |
| NE | 0.2047 | 0.2180 | 0.0612 |
| NW | 0.2060 | 0.2192 | 0.0602 |
| SW | 0.1982 | 0.2151 | 0.0787 |
| PW | 0.1761 | 0.1990 | 0.1152 |
| ESM | 0.1740 | 0.1890 | 0.0797 |
| WSM | 0.1797 | 0.1892 | 0.0502 |

### Historical population size changes per *P. tremuloides* lineage

All genetic lineages displayed a historical bottleneck pattern starting between 0.5-2 million years ago (mya) (**Fig. 6**). In detail, the ESM lineage showed a median effective population size (*N_e_*) decline starting around 1.5 mya, that reached its lowest value (*N_e_* ≈ 260,000) around 0.5 mya after which it bounced back to an effective population size of 3.3 million around 400 kya and has remained stable ever since.

**Figure 6.**
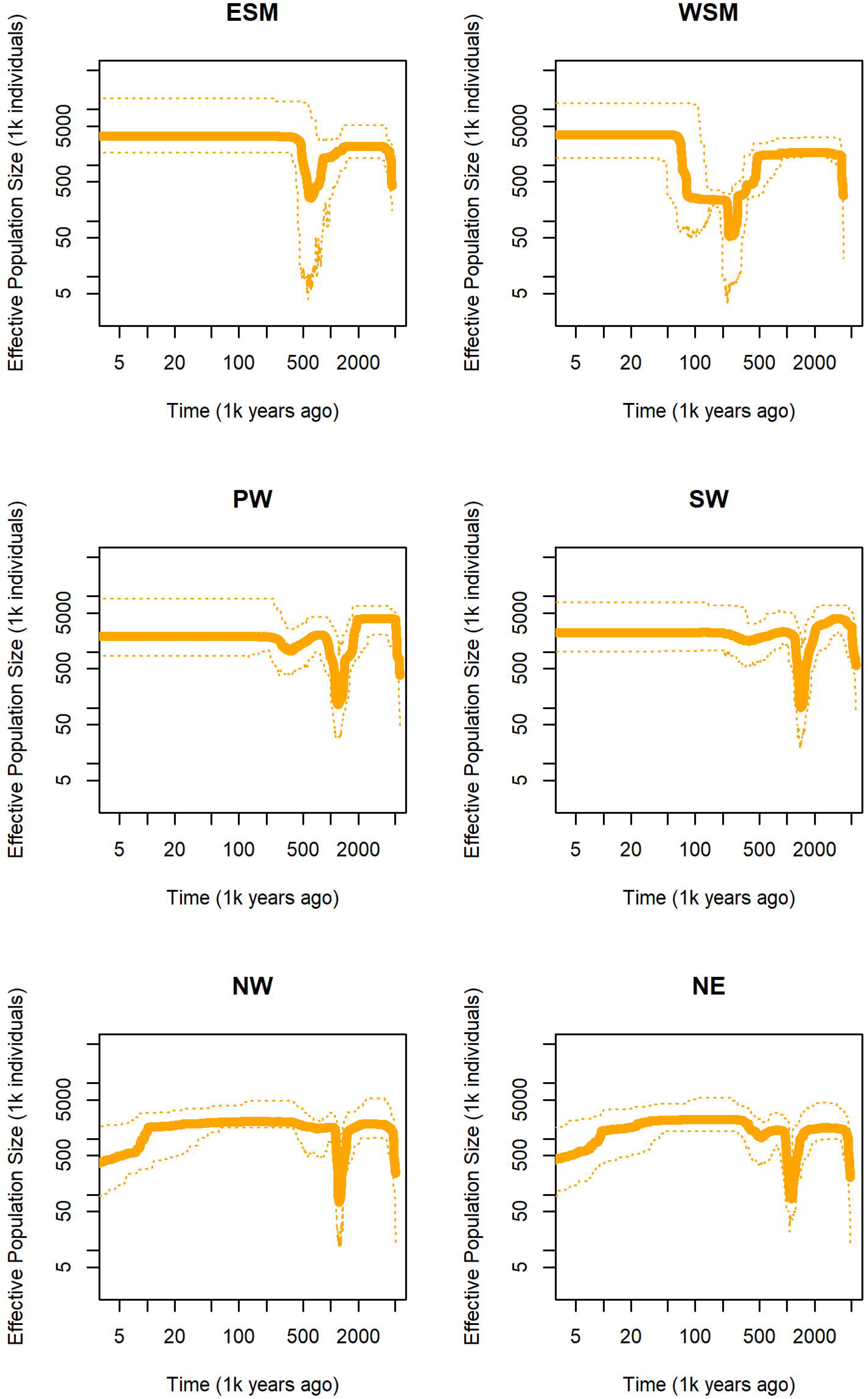
Stairway Plot 2 results showing aspen lineages’ demographic trajectories. ESM: Sierra Madre Oriental, WSM: Sierra Madre Occidental, PW: Pacific West, SW: southwestern US, NW: northwest North America and NE: northeast North America. Dashed lines indicate the 95% confidence intervals.

The WSM lineage displayed a bottleneck starting around 1 mya, leading to its lowest value in *N_e_* around 0.25 mya (*N_e_* ≈ 53,000), after which *N_e_* increased to around 240,000 and remained stable until around 80 kya. Hereafter, *N_e_*abruptly increased to 3.5 million around 65 kya and has remained stable until the present day.

The PW lineage displayed an *N_e_* decline starting around 2 mya, reaching its low around 1.3 mya (*N_e_* ≈ 114,000), and followed by an *N_e_* increase nearing 2 million around 0.8 mya. Hereafter, the *N_e_* steadily decreased but only dropped to 1 million around 350 kya. Hereafter an increase in *N_e_* is observed up to 1.9 million around 300 kya and has remained stable until now.

The SW lineage displayed a pattern like the PW lineage, with an *N_e_* decline starting around 2 mya, reaching a low *N_e_* of 102,500 around 1.5 mya. Hereafter, we observed an increase in *N_e_* of up to 2.3 million around 1.2 mya, followed by a slow decline around 400 kya and a subsequent increase (*N_e_* ≈ 2.2 million) around 200 kya and remaining stable until now.

The NE and NW lineages displayed a highly similar historical *N_e_*pattern, with an *N_e_* decline starting around 1 mya reaching a low of 85 k, and 72 k around 1.4 mya. Hereafter, both lineages increased until 0.9 mya to, respectively, an approximate *N_e_* of 1.4 million and 1.6 million for NE and NW. A small decline followed, while more pronounced in the NE lineage, followed by an increase and stable *N_e_* up until approximately 80 kya. Hereafter, both lineages showed a decline until the present day to an *N_e_* of 863 and 4,218 for NE and NW, respectively.

### Sex distribution in *P. tremuloides* over its entire range

We identified over the entire sample size of 1734 individuals 1021 clonal groups, ranging in size from 1 to 30 clonemates. We combined our previous ploidy identification results (Goessen et al., 2025) with our current results on Texas samples, resulting in a total of 1,474 diploid, 170 triploid, and only 20 ambiguous individuals. Among all tested individuals, we identified 853 females and 997 males (including the Texas individuals) (**Dataset S3**).

After combining our ploidy, clonality and sex determination results (including the removal of missing data for either of these three trait aspects), we obtained 1,363 individuals, and subsequently after considering clonemates, we obtained a total of 724 genetically unique individuals, involving 410 males and 314 females (**Dataset S4**). Taking ploidy into account, these consisted of 10 triploid females, 37 triploid males, 304 diploid females and 373 diploid males (**Fig. 7A-B**, **Table 3**). Male individuals exhibited a substantially higher occurrence of triploidy (37/410, 9.02%) compared to females (10/314, 3.18%). The chi-square test between males and females regardless of ploidy, resulted in an X-squared test statistic of 12.729, and p-value of 0.0004, indicating that there is no equal proportion between males and females, specifically indicating a male-bias. Further analysis of range-wide sex distribution by geographic region revealed a male bias in several genetic lineages. The NW lineage showed the strongest male overrepresentation, with 115 males and 68 females, followed by the NE lineage with 129 males and 104 females. The WSM lineage included 120 males and 99 females, and the ESM lineage had 19 males and 15 females. In contrast, the PW lineage had a near-equal sex ratio (14 males, 15 females), as did the SW lineage (13 males, 13 females) (**Fig. S4A, Table 3, Dataset S4**).

**Figure 7.**
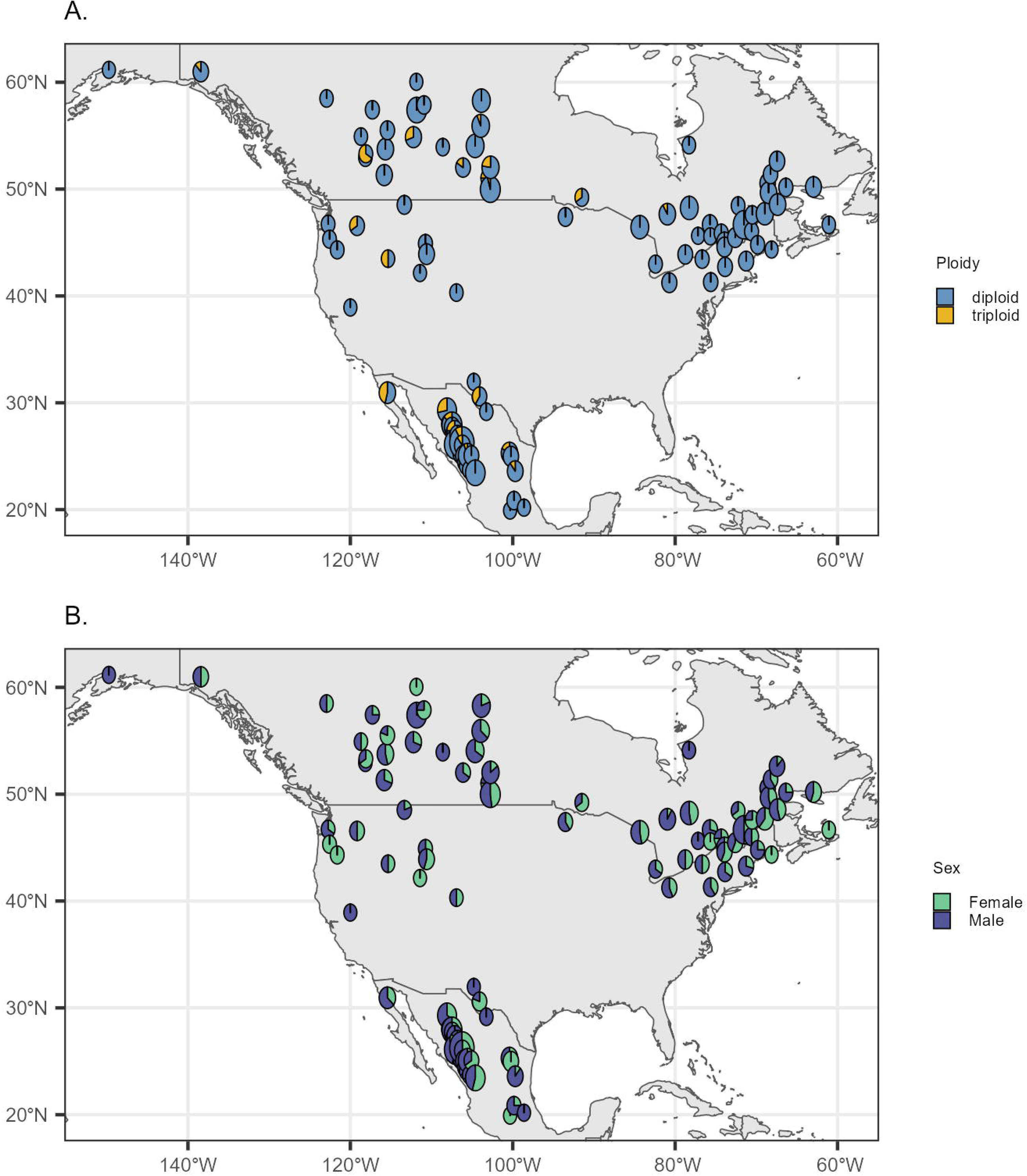
Distribution of ploidy and sex summarized per population after clone-correction (i.e. removal of clonemates) across the sampled aspen distribution range. **A.** Ploidy distribution per population. **B.** Sex distribution per population. See **Table 3** for exact numbers of sex and ploidy per genetic lineage.

**Table 3.** Distribution of sex and ploidy in aspen per genetic lineage related to geographic regions (K=6) across the sampled aspen distribution range. ESM: Sierra Madre Oriental, WSM: Sierra Madre Occidental, PW: Pacific West, SW: southwestern US, NW: northwest North America and NE: northeast North America. For a visual representation of sex and ploidy across aspen populations, see **Fig. 7**.

| Region/Lineage |  |  |  |  |  |  |  |  |
| --- | --- | --- | --- | --- | --- | --- | --- | --- |
| Sex | Ploidy | NE | NW | PW | SW | WSM | ESM | Total |
| Male | Diploid | 128 | 103 | 11 | 6 | 108 | 17 | 373 |
|  | Triploid | 1 | 12 | 3 | 7 | 12 | 2 | 37 |
| Female | Diploid | 104 | 66 | 15 | 11 | 94 | 14 | 304 |
|  | Triploid | 0 | 2 | 0 | 2 | 5 | 1 | 10 |
|  | Total | 233 | 183 | 29 | 26 | 219 | 34 |  |

The Chi-square test on tree sex within diploids found a X^2^ test statistic of 7.0325 and p-value of 0.008, again confirming that the sex ratio in diploids does not conform to a 50:50 ratio. When assessing sex within triploids, the X-squared test statistic was 15.511 with a p-value of 0.00008, indicating that there is a stronger male-bias present among triploids. Most triploid males were present in the WSM lineage (12 triploid males and 5 triploid females), followed by the NW lineage with 12 triploid males and 2 triploid females, and the SW lineage with 7 triploids males and 2 triploid females. The GLM with sex as dependent and elevation as independent variable revealed no significant effect (p-value = 0.508) of elevation on sex occurrence over the entire distribution range (**Table S1**). When we assessed the GLM per genetic lineage, we identified a weak significant relationship (p-value of 0.048) between more males at higher elevations (effect of 0.0091) for the SW lineage; however, this lineage also has the lowest sample size of 26 (**Table S2**). Other genetic lineages do not display any significant effect (**Table S2**).

## Discussion

This study provided new insights into the genetic composition, clonality, ploidy, and sex distribution within *Populus tremuloides*, with a particular focus on previously uncharacterized populations in Texas. By integrating these new data within our prior range-wide genomic dataset, we revealed that Texan aspen stands have been shaped by substantial clonality, include triploid and diploid individuals, mixed sex, and show strong genetic affiliation with southwestern and Mexican genetic lineages, respectively. These findings extend previous microsatellite-based assessments (Nunneley et al. 2014) and also further confirm occurrences of triploidy within arid environments (Mock et al. 2012; Goessen et al. 2022). Environmental aridity also predicted cytotype distribution and polyploidy occurrence in temperate annual grass *Brachypodium distachyon* in the Iberian Peninsula (Manzaneda et al., 2012). Moreover, our range-wide analyses in aspen highlighted a significant male bias in sex ratios, particularly among triploids. Elevation did not significantly affect sex distribution across the species’ range or within genetic lineages. While the SW lineage showed a weakly significant effect, we interpret this finding cautiously given the limited sample size from this region.

Our findings confirm that many of the Texan aspen stands are dominated by clonal groups, consistent with previous reports of widespread clonality in isolated southern populations (Goessen et al. 2022, 2025; Nunneley et al. 2014). In particular, we found that individuals from stands DMP02, DMP03, and DMP04, that are located at Laura’s Rock, are genetically near identical and share the same sex, strongly suggesting that they form a single large female clone. This supports earlier microsatellite-based conclusions by Nunneley et al. (2014), who identified these same stands as genetically similar. Similarly, individuals from DMP05 and DMP06, corresponding to the BT and PWS stands in Nunneley et al. (2014), were previously considered highly related but not clonal. Our SH-index analysis under both loose and strict filtering thresholds indicates SH-values near or above 0.9 between most individuals in these stands. Their shared male sex, triploid assignment and spatial proximity further support the hypothesis that these stands may form a single male triploid clone.

We found a significant male bias in sex ratios at the species level, that was even more pronounced among triploids. While we cannot say for certain what the causes are for such male bias, we can hypothesize several reasons. Potentially, males produce more unreduced gametes than females, causing more male triploids to arise. A higher production of male unreduced gametes has for instance been observed in certain crosses between cultivated sunflower and wild accessions (Liu et al., 2017). Under environmental stress, it is possible that males may have an advantage due to sex-differential reproductive costs compared to females (Barrett et al., 2010), i.e. the production of pollen vs seeds, and other secondary sex characteristics, such as morphological and physiological traits (Liu et al., 2021). This advantage may be more pronounced in triploids considering their lower assumed fertility.

The incorporation of Texas individuals into our ADMIXTURE and phylogenetic analyses found that the stands from Guadalupe Mountains and Davis Mountains mostly belong to the southwestern (SW) genetic lineage. While individuals from Big Bend National Park clustered with the WSM lineage in Mexico. Previous research identified that the earliest split among genetic lineages of quaking aspen occurred around 800 kya in the southern part of the species’ range, based on DIY-ABC-RF analyses (Goessen et al. 2025). Since these Texan individuals appear to bridge these genetic lines phylogenetically, they underscore the role of the Trans-Pecos region as a transitional zone that may have shaped ancient southern divergence. The studied sky island mountain ranges in this Trans-Pecos region are separated by lowland Chihuahuan desert vegetation (Muldavin, 2002; Nadeau et al., 2014). Other tree species that share this habitat include (among others) Douglas-fir (although absent in Davis Mountains), Ponderosa pine, and several oak species (Elliott, 2014). However, to our knowledge, barely any population genomics studies have been performed on these species that cover the Trans-Pecos region. One study on several species in the subsection of *Ponderosaea* assessed mitochondrial haplotypes, covering two locations in the Trans-Pecos region (Willyard et al., 2021). They found that one haplotype included individuals from northern Mexico, southern California, Guadalupe and Davis Mountains, as well as mountains in the Mojava Desert in southern Nevada. This haplotype was found to be most similar to the most eastern US group and haplotypes only present in Mexico (Willyard et al., 2021). A study on Douglas-fir covered populations of southern New Mexico (US) located near the Trans-Pecos region, and southern Mexico, and found that these populations were part of the same genetic lineage, however, due to the sparse sampling in Mexico (2 populations only) this result may be biased (Peláez et al., 2024).

All lineages displayed a historical bottleneck around 1-2 million years ago, potentially corresponding to global cooling. Against our expectations, the NE and NW lineages displayed a slight decline in *N*_e_ since around 80 kya, which is in contrast with results produced by fastsimcoal2 in previous studies, which observed a population expansion between 50 and 80 kya in these lineages (Wang et al., 2016; Goessen et al., 2025). This finding could be interpreted as methodological differences, whereby Stairway Plot 2 estimates continuous *N*_e_ change without specifying a model, while fastsimcoal2 requires explicit demographic scenarios. The other four genetic lineages displayed a stable *N*_e_ scenario for after the bottleneck recovery since at least 500 kya, while a recent bottleneck was previously found using fastsimcoal2 in the southern genetic lineage (Goessen et al. 2025). Again, it is known that methods that infer *N_e_* continuum over time, such as Stairway Plot 2, can be less accurate for recent time periods and might thus miss signals in *N_e_* changes (Patton et al., 2019; Nadachowska Brzyska et al., 2022). While Stairway Plot 2 is used for reduced representation sequencing data, whole genome sequencing data with appropriate demographic inference tools in future may provide conclusive outcomes with higher resolution (Sandercock et al., 2022),. These types of tools can further substantiate evidence of the distinct demographic histories of identified genetic lineages, which can be recognized as separate conservation units for management purposes.

## Conclusion

Taken together, our findings provide a comprehensive genetic and demographic portrait of *Populus tremuloides* populations at the southern margin of the species’ range. By integrating Texan aspen stands into a range-wide genomic framework, we uncovered their genetic links to southwestern and Mexican lineages, confirmed high levels of clonality and triploidy in arid environments, and detected notable biases in sex ratios, particularly among triploids. These results refine our understanding of how marginal, isolated populations are structured and maintained, and highlight how clonality, polyploidy, and sex interplay in shaping the biology of this widespread species.

## Acknowledgements

We thank U.S. Parks for allowing us to sample Guadalupe Mountains National Park and Big Bend National Park, and we thank the Nature Conservancy for allowing us to sample at Davis Mountains Preserve. We acknowledge the Discovery grant from the Natural Sciences and Engineering Research Council of Canada (RGPIN/04748-2017) to Ilga Porth as well as the GRDI funding to Nathalie Isabel. We thank Digital Alliance Canada for using their computational resources. We thank Melanie Zacharias for helping with DNA concentration testing. We thank Katrin Groppe, Thünen Institute of Forest Genetics, Germany, for her previous work on aspen sexing. Lastly, we thank everyone involved in the range-wide aspen collection.

## Author contribution

IP conceived the overall study and provided funding for the study. RG, IP, CD contributed to experimental design and interpreted the data. RG performed DNA extractions, all genomic analyses and wrote the manuscript. IP, CD revised the manuscript. RG, LT, NI, IP performed sex determination work. CD, ER, JN were involved in Texas sample collection and NI in Texas samples genotyping.

## Data availability

Raw sequencing data are deposited in the Sequence Read Archive (SRA) under BioProject PRJNA809939 and will be released upon publication.

