## Supplements for "Male-biased triploidy in *Populus tremuloides* and genomic structure of its southern relict populations"

The following Supporting Information is available for this article:

**Figure S1.** Ploidy assessment with *Fastploidy* for Texan samples.

**Figure S2.** Admixture cross-validation for selection of K clusters for the two datasets of sampled aspen (including Texas).

**Figure S3.** Pairwise $F_{ST}$ between genetic lineages for the diploid dataset of sampled aspen (including Texas).

**Figure S4 A-C.** Distribution of sex over geographic regions across genetic lineages and elevation for sampled aspen (including Texas).

**Table S1.** Results of generalized linear model with elevation as independent variable and sex as dependent variable across sampled aspen (including Texas).

**Table S2.** Results of generalized linear model with elevation as independent variable and sex as dependent variable per genetic lineage (including Texas).

**Dataset S1.** Overview of sampling data and assignment of genetic lineages across sampled aspen (including Texas).

**Dataset S2.** Pairwise shared heterozygosity index (SH-index) for loose and strict filtered dataset.

**Dataset S3.** Overview of combined sex, ploidy and clonality assignment prior to filtering across sampled aspen (including Texas).

**Dataset S4.** Overview of combined sex, ploidy and clonality assignment after filtering across sampled aspen (including Texas).

### Supplemental figures

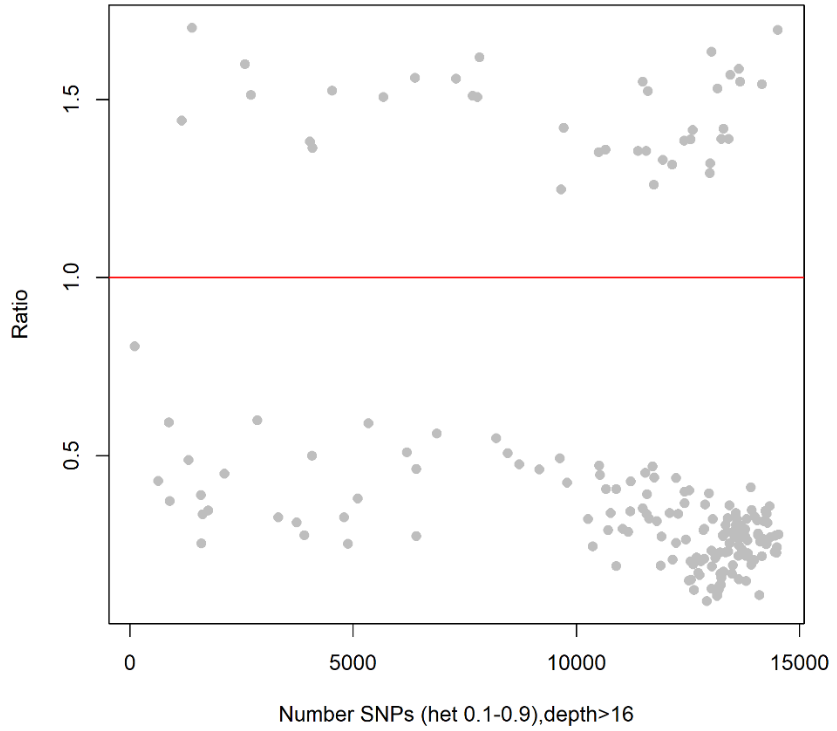


**Figure S1. Ploidy assessment with *Fastploidy* for Texan samples.** Samples above the red line are assigned triploids.

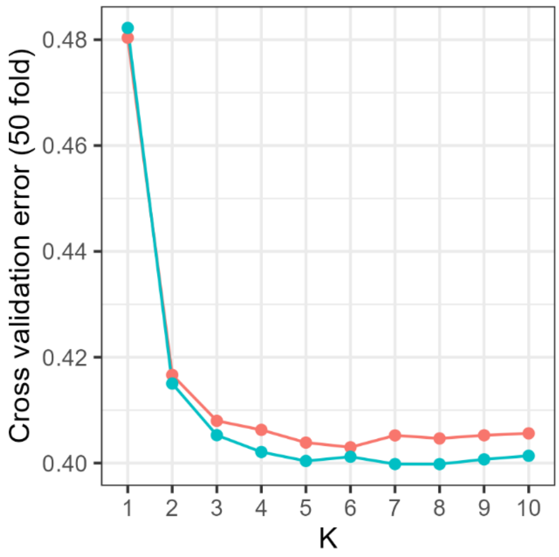


**Figure S2. Admixture cross-validation for selection of K clusters for the two datasets of sampled aspen (including Texas).** The best K is K=5 for the mixed ploidy (blue line) and K=6 for the diploid (red line) dataset.

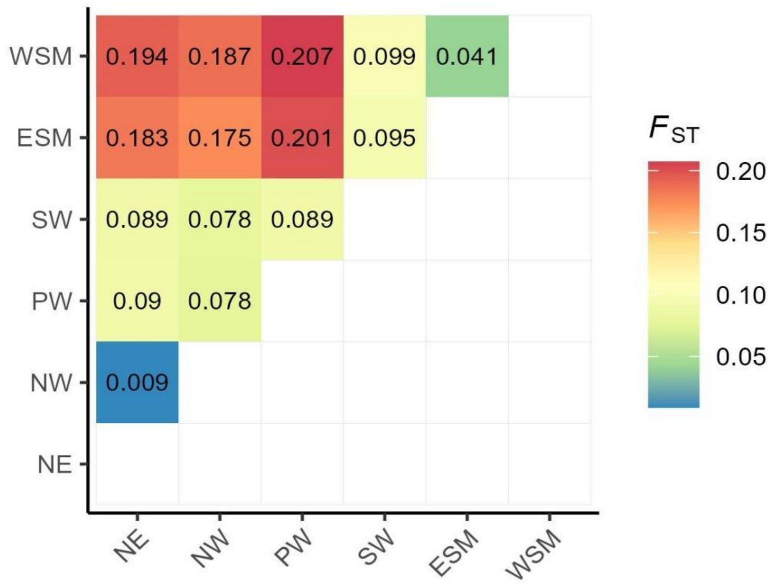


**Figure S3**. **Pairwise $F_{ST}$ between genetic lineages for the diploid dataset of sampled aspen (including Texas).** ESM: Sierra Madre Oriental, WSM: Sierra Madre Occidental, PW: Pacific West, SW: southwestern US, NW: northwest North America and NE: northeast North America.

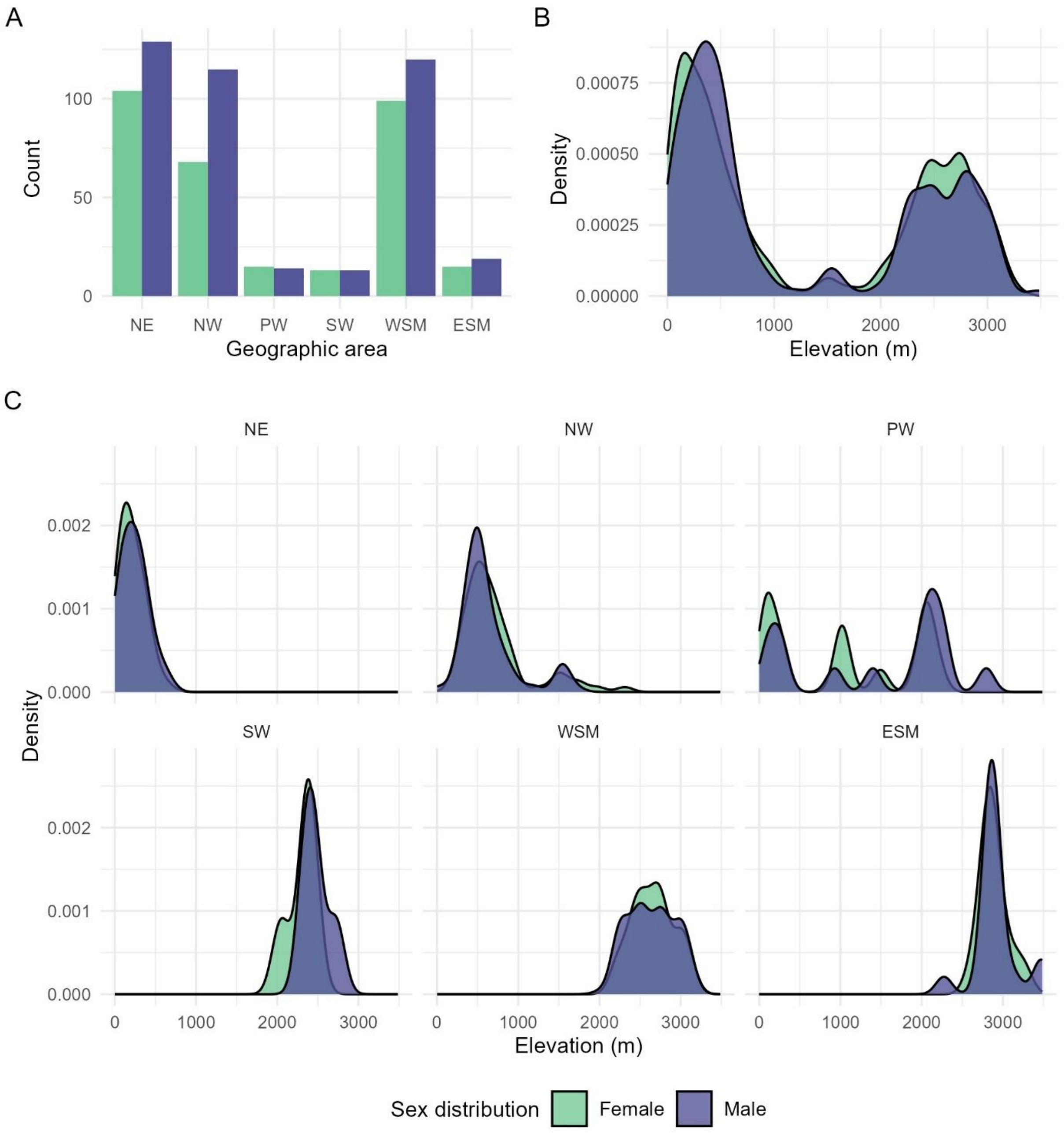


**Figure S4 A-C. Distribution of sex over geographic regions across genetic lineages and elevation for sampled aspen (including Texas)**. A. Count of males and females per geographic region. B. Density plot of sex across elevations over the entire distribution range. C. Density plot of sex across elevations for individual geographic regions**.** Density is calculated over 100-meter windows. ESM: Sierra Madre Oriental, WSM: Sierra Madre Occidental, PW: Pacific West, SW: southwestern US, NW: northwest North America and NE: northeast North America.

**Supplemental tables**

**Table S1. Results of generalized linear model with elevation as independent variable and sex as dependent variable across sampled aspen (including Texas).** Results indicate no significant effect of elevation on sex occurrence across the sampled aspen distribution range.

| | Estimate | Std. Error | z value | Pr(>\|z\|) |
|---|---|---|---|---|
| Intercept | 0.32432 | 0.115031 | 2.819426 | 0.004811 |
| elevation | -4.4E-05 | 6.6E-05 | -0.6618 | 0.508102 |

**Table S2. Results of generalized linear model with elevation as independent variable and sex as dependent variable per genetic lineage (including Texas).** Results indicate a significant effect for the SW lineage, but no significant effect for the other lineages.

| | Estimate | Std. Error | zvalue | Pr(>\|z\|) | Cluster | Size |
|---|---|---|---|---|---|---|
| Intercept | -0.0880 | 0.2321 | -0.3793 | 0.7045 | NE | 233 |
| elevation | 0.0014 | 0.0009 | 1.5753 | 0.1152 | | |
| Intercept | 0.8848 | 0.3070 | 2.8820 | 0.0040 | NW | 183 |
| elevation | -0.0005 | 0.0004 | -1.3640 | 0.1726 | | |
| Intercept | -0.7624 | 0.6730 | -1.1328 | 0.2573 | PW | 29 |
| elevation | 0.0005 | 0.0004 | 1.2641 | 0.2062 | | |
| Intercept | -21.6601 | 11.0123 | -1.9669 | 0.0492 | SW | 26 |
| elevation | 0.0091 | 0.0046 | 1.9695 | 0.0489 | | |
| Intercept | 0.9874 | 1.3294 | 0.7427 | 0.4576 | WSM | 219 |
| elevation | -0.0003 | 0.0005 | -0.6014 | 0.5476 | | |
| Intercept | -1.3664 | 4.7552 | -0.2874 | 0.7738 | ESM | 34 |
| elevation | 0.0006 | 0.0016 | 0.3378 | 0.7355 | | |

**Supplemental datasets**

**Dataset S1. Overview of sampling data and assignment of genetic lineages across sampled aspen (including Texas).**

Will be uploaded as excel sheet.

**Dataset S2. Pairwise shared heterozygosity index (SH-index) for loose and strict filtered dataset.**

Will be uploaded as excel sheet.

**Dataset S3. Overview of combined sex, ploidy and clonality assignment prior to filtering across sampled aspen (including Texas).**

Will be uploaded as excel sheet.

**Dataset S4. Overview of combined sex, ploidy and clonality assignment after filtering across sampled aspen (including Texas).**

Will be uploaded as excel sheet.
